# Time to budbreak is not enough: cold hardiness evaluation is necessary in dormancy and spring phenology studies

**DOI:** 10.1101/2022.09.15.508138

**Authors:** Michael G. North, Al P. Kovaleski

## Abstract

Dormancy of buds is an important phase in the life cycle of perennial plants growing in environments where unsuitable growth conditions occur seasonally. In regions where low temperature defines these unsuitable conditions, the attainment of cold hardiness is also required to survive. The end of the dormant period culminates in budbreak and flower emergence, or spring phenology, one of the most appreciated and studied phenological events. Despite this, we have a limited physiological and molecular understanding of dormancy, which has negatively affected our ability to model budbreak. Here we highlight the importance of including cold hardiness in studies that typically only characterize time to budbreak. We show how different temperature treatments may lead to increases in cold hardiness, and by doing so also (inadvertently) increase time to budbreak. Therefore, erroneous interpretations of data may occur by not phenotyping cold hardiness. Changes in cold hardiness were very likely present in previous experiments to study dormancy, especially when those included below freezing temperature treatments. Separating the effects between chilling accumulation and cold acclimation in future studies will be essential for increasing our understanding of dormancy and spring phenology in plants.

## Main text

Dormancy, along with development of cold hardiness in tissues, allows plants to survive unsuitable growing conditions during winter and precisely time budbreak upon return of suitable temperatures in spring. Chilling accumulation – the exposure to low temperatures for a period of time – promotes the transition from the warm temperature non-responsive phase to the warm temperature-responsive phase due to ontogenetic changes within buds (often referred to as endo-to ecodormancy transition – see Lang et al., 1987 for definitions). The molecular and physiological basis for dormancy and its transitions remain only partially understood (Cooke et al., 2012; Yamane et al., 2021). In turn modeling chilling accumulation across different regions (Luedeling and Brown, 2011) and modeling time to budbreak in spring (spring phenology) (Melaas et al., 2016; Wang et al., 2020; Zohner et al., 2020) present a linked challenge.

Here we show that the phenotype of time to budbreak, which has been used in the vast majority of experiments for over a century, only tells part of the story. This shortcoming has limited advances in our understanding of dormancy from a mechanistic standpoint, and related aspects such as the development of accurate chilling accumulation and spring phenology models, which is an important component of Earth system models (Richardson et al., 2012; Chen et al., 2016). We argue that evaluation of cold hardiness and deacclimation is necessary to accurately interpret budbreak as a phenotype for dormancy completion. To do so, we present here a combination of original data along with re-analysis of previously published data from Kovaleski (2022).

The first presumed report of low temperature exposure (chilling) as a requirement for proper budbreak of temperate species once exposed to warm temperatures (forcing) is over two centuries old (Knight, 1801). Chilling-forcing experiments have been the standard approach to study dormancy for at least 100 years (Coville, 1920). In these experiments, plants or cuttings are subjected to low temperatures for varying durations (chilling treatments), either naturally (field) or artificially (low temperature chambers), and then transferred to forcing conditions to monitor regrowth. The typical metrics recorded in these assays are based on visual observation of percent budbreak and/or time to budbreak (Londo and Johnson, 2014; Alvarez et al., 2018). Longer duration of chilling treatments correlates with higher percent budbreak and shorter time to budbreak (i.e., negative correlation between chilling accumulation and heat requirement under forcing). However, in most studies the temperature treatments applied as chilling are also inadvertently affecting other physiological aspects in the buds beyond dormancy progression, including cold hardiness.

While artificial chilling treatments are often described as constant, positive temperatures (e.g., 1.5 ºC and 4 ºC within Flynn and Wolkovich (2018)), more and more studies have included negative temperatures to study their effect on chilling accumulation (–3 ºC, –5 ºC and –8 ºC in Cragin et al., 2017; –2 ºC in Baumgarten et al., 2020). However, these experiments have not included evaluations of cold hardiness in response to chilling. The combined effects of chilling and cold hardiness on time to budbreak have only been studied in field conditions, although this has now been done in many species, both of fruit crops, such as grapevines (Kovaleski et al., 2018; Kovaleski 2022; North et al., 2022) and apricot (Kovaleski, 2022), and other ornamental and forest species (Lenz et al., 2013; Vitra et al., 2017; Kovaleski, 2022). However, artificial chilling experiments are key to better understand effects of particular temperatures in providing chilling, as field conditions are too variable for this.

Using grapevine cuttings of five different cultivars (all *Vitis* interspecific hybrids), we supplied chilling using three different treatments: constant (5 ºC), fluctuating (–3.5 °C, 6.5 ºC, for 7h, 17h intervals daily), and field collected cuttings in Madison, WI, USA. When evaluating cold hardiness of the buds, we observed that the fluctuating treatment elicited a greater gain in cold hardiness over time compared to constant, while both were surpassed by buds subjected to much colder temperatures in the field (Fig. **1a**). After 2.5 months under treatments, some cuttings from constant and fluctuating treatments were reciprocally exchanged. Cold hardiness was again evaluated one month after the reciprocal exchange: field buds were still the most cold hardy, followed by all treatments which had been at any point exposed to fluctuating conditions, while buds that remained in the constant temperature treatment were the least cold hardy (Fig. **1b**, bottom). Cuttings from the same treatments, when placed under forcing conditions (22 ºC, 16h/8h day/night) for time to budbreak evaluation, demonstrate a similar, but opposite distribution: field collected cuttings take the longest to break bud, whereas constant temperature treated buds take the least amount of time (Fig. **1b**, top). Based on the observations of time to budbreak alone, the interpretation would be that the constant temperature treatment was the most effective in supplying chilling to buds, leading to shorter time to budbreak compared to fluctuating and field. However, even though exposure to all treatments reduced time to bud break by providing considerable chilling, the chilling effects in fluctuating and field treatments were diminished by the elongation of time to budbreak attributable to pronounced gains in cold hardiness.

**Figure 1.**
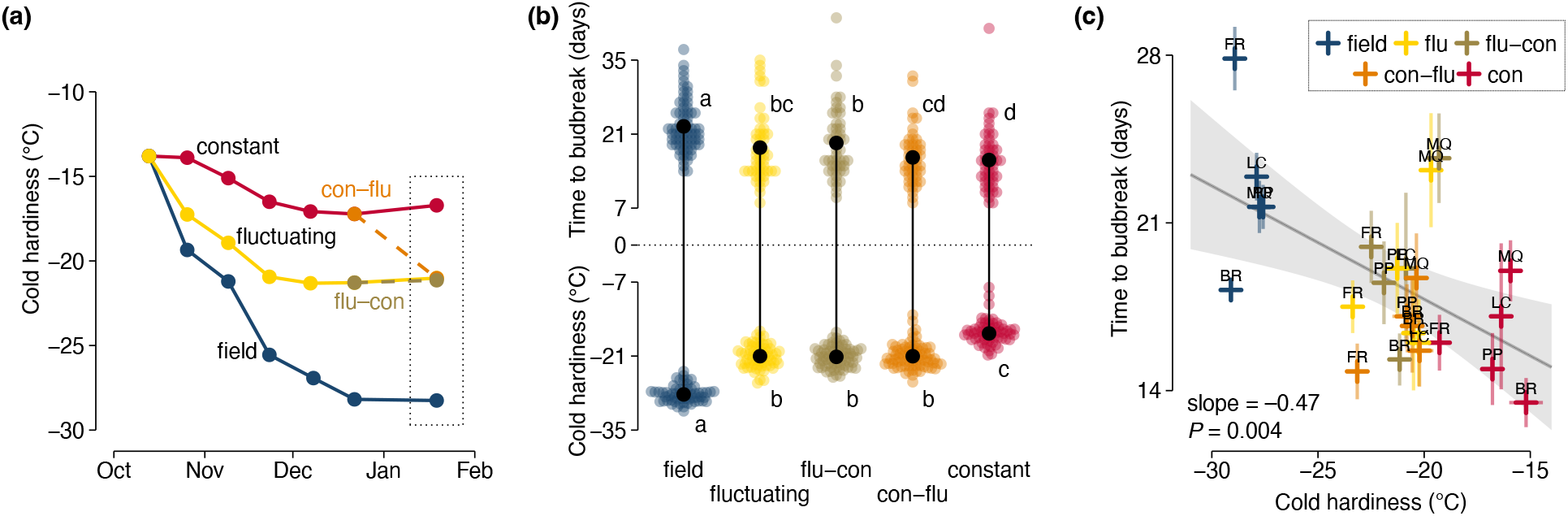
Cold hardiness and time to budbreak relations of grapevine buds in response to different chilling treatments. Cuttings of five *Vitis* interspecific hybrid cultivars (‘Brianna’ – BR, ‘Frontenac’ – FR, ‘La Crescent’ – LC, ‘Marquette’ – MQ, ‘Petite Pearl’ – PP) were exposed to three different chilling treatments: constant temperature (5 ºC), fluctuating temperature (–3.5 °C, 6.5 ºC, for 7h, 17h intervals daily), and field temperatures (in fall and winter of 2021-2022, Madison, WI, USA). After 2.5 months under treatment, cuttings from the artificial chilling treatments were reciprocally exchanged. **(a)** Cold hardiness of all original treatments was measured using 15 buds in bi-weekly intervals until the exchange point, and a final cold hardiness measurement was performed one month after the exchange. **(b)** Pairwise comparisons of cold hardiness and time to budbreak under forcing conditions (22 ºC, 16h/8h day/night) for all cultivars combined using Fishers LSD at α = 0.05. **(c)** Linear model showing a relationship between time to budbreak and cold hardiness of individual cultivar samples. Standard error of observations is illustrated as semi-transparent extensions from points horizontally (time to budbreak) and vertically (cold hardiness). See experiment description in SI Materials and Methods.

The relationship between time to budbreak and cold hardiness provides us with additional information. A slope of about –0.5 day ºC^-1^ is observed when looking at the relationship of cold hardiness to time to budbreak (Fig. **1c**). This means that for every two additional degrees Celsius of cold hardiness, buds will take an additional day to break bud. The inverse of this slope is also useful: if we consider budbreak occurs at the end of the cold hardiness loss period, we can estimate a deacclimation rate of approximately 2 ºC day^-1^ based on these data (approximately the maximum deacclimation rate reported by North et al. (2022) for the same cultivars). Here this is measured at high levels of chill accumulation, after 3.5 months under chilling treatments, where deacclimation responses are likely maximized. At low chilling accumulation, the slope in Fig. **1c** would presumably be higher due to lower deacclimation rates (Kovaleski et al., 2018; Kovaleski, 2022; North et al., 2022).

This effect is not confined to *Vitis* spp.: buds of many other species, both angiosperms and gymnosperms, deciduous and evergreen, gain cold hardiness during exposure to low temperatures, particularly when negative temperatures are included in treatments [Fig. **2a**; see also hardening treatment in Vitra et al. (2017)]. The relevance of cold hardiness gains in relation to time to budbreak depends on how much each species responds (Fig. **2b**) (where higher gains will have a greater effect) and how quickly any given species loses cold hardiness (see Box 1).

**Figure 2.**
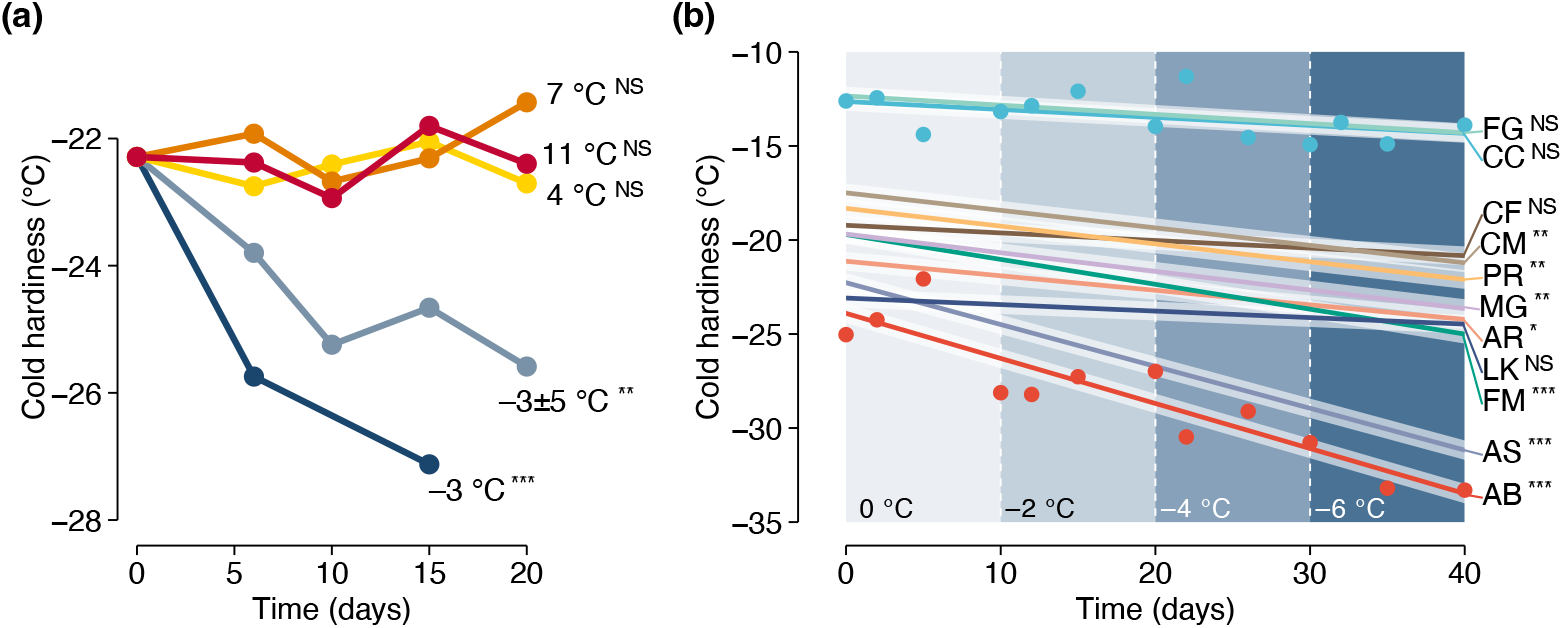
Cold hardiness changes in response to different temperature treatments for different ornamental and forest species. Changes in cold hardiness were analyzed using linear models in response to time. **(a)** Combined effect of temperature treatments on bud cold hardiness of *Acer platanoides, A. rubrum, A. saccharum, Cornus mas, Forsythia* ‘Meadowlark’, *Larix kaempferi, Metasequoia glyptostroboides, Picea abies, Prunus armeniaca*. Temperature treatments were constant –3 ºC, 4 ºC, 7 ºC, 11 ºC and fluctuating (−8 ºC, −3 ºC, 2ºC, −3 ºC for 6h intervals each: “−3±5 ºC”). **(b)** Cuttings of eleven species were exposed to decreasing temperatures in –2 ºC steps every 10 days from 0 ºC to –6 ºC. Linear responses are shown for all species, along with data points for two species: *Cercis canadensis* (CC) and *Abies balsamea* (AB). Other species include: FG – *Fagus grandifolia*; CF – *Cornus florida*; CM – *Cornus mas*; PR – *Prunus armeniaca*; MG – *Metasequoia glyptostroboides*; AR – *Acer rubrum*; LK – *Larix kaempferi*; AS – *Acer saccharum*. Asterisks indicate level of significance of slopes for linear models of cold hardiness in response to time in **(a)** and **(b)**: ^NS^not significant; ^*^*P* ≤ 0.05; ^**^*P* ≤ 0.01; ^***^*P* ≤ 0.001. See experiment description in SI Materials and Methods.

It is clear that low temperatures – particularly negative temperatures – in chilling treatments can lead to increases in cold hardiness (Fig. **1a**,**b** and Fig. **2a**,**b**), and by doing so can increase time to budbreak (Fig. **1b**,**c**). However, previous studies have not taken into consideration the effect of cold hardiness gains increasing time to budbreak. For example, Cragin and colleagues (2017) showed negative temperatures to contribute differently in the chilling accumulation of two grapevine genotypes: –3 ºC was more effective than 0 ºC and 3 ºC for ‘Chardonnay’, but the opposite for ‘Cabernet Sauvignon’. It is possible that the increases in rate of deacclimation elicited by chilling at the negative temperatures in ‘Cabernet Sauvignon’, which should lead to faster budbreak, was balanced by gains in cold hardiness, leading to a perceived delay in in time to budbreak (e.g., Box **1d**).

Baumgarten and colleagues (2021) showed that a high but sub-freezing temperature (–2 ºC) does contribute to chilling of many forest species, though at different magnitudes. Notably, negative temperature treatments seemed to be more effective than many other low above freezing temperature treatments – consistent with findings for ‘Chardonnay’ by Cragin et al. (2017). Given the likely effect of the negative temperature eliciting greater gains in cold hardiness (e.g., Fig. 2), it is possible that the effect of this treatment in providing chilling is underestimated there: if we account for the additional days taken to break bud because of the greater cold hardiness of buds, it may be that such temperatures are even more effective in providing chilling than what was estimated. Similarly, Rinne and colleagues (1997) applied short-term freezing treatments (–8 °C, –16 °C, –24 °C, –32 °C) to *Betula pendula* seedlings during the dormant period. They observed a slight increase in days to budbreak (from four to eight weeks) before subsequent declines (from eight to twelve weeks). This could be explained by the simultaneous but competing effects between acclimation, which leads to increases in time to budbreak, and chilling accumulation, which leads to decreases in time to budbreak. In these non-exhaustive examples we speculate low temperatures are not only promoting dormancy transitions but are also promoting acclimation. However, these effects cannot be separated without cold hardiness measurements. Therefore, including cold hardiness measurements in future studies could clarify our understanding of the range of temperatures promoting chilling and lead to improved chilling and phenology models.

Most phenological models only use combinations of chilling and forcing as temperature effects in their predictions (Wolkovich et al., 2012; Melaas et al., 2016; Vitasse et al., 2018; Ettinger et al., 2020; Zohner et al., 2020). Within the work of Melaas and colleagues (2016), it is interesting to note that the error in spring onset predictions follows a clear climatic gradient for many species, possibly indicating changes in cold hardiness along this gradient [though other genotypic differences can also play a role (Thibault et al., 2020)]. Recently, Wang and colleagues (2020) attempted to include a term for cold hardiness, but this resulted in no improvement over simpler models. However, they only compared “low” and “high” latitudes, dividing their dataset at 50.65º N. By doing so, the high latitude combined data from areas with much milder climates, such as the British Isles, with data from much colder areas, such as the Nordic countries. While a division based on minimum observed temperatures might be a more sensible approach in modeling, it would possibly still not be enough given the dynamic nature of cold hardiness. It is also important to consider the duration of cold exposure based on incremental cold hardiness gains over time in artificial treatments (Fig. 1 and Fig. 2), something that is often acknowledged in field cold hardiness models (Aniśko et al., 1994; Ferguson et al., 2011; Ferguson et al., 2014).

Cold hardiness and dormancy are thus intrinsically connected. In particular, dormancy establishment (or at least growth cessation) precludes significant acclimation (Tanino et al., 2010). Similar low temperatures promote gains in cold hardiness and chilling accumulation (Fig. **3a**). Increasing chill accumulation leads to increases in rates of cold deacclimation (Kovaleski et al., 2018; North et al., 2022; Kovaleski, 2022). Given these overlaps, could chilling and acclimation both be part of the same process? A correlation between both has been previously suggested (Wolf and Cook, 1992; Cragin et al., 2017). However, here we argue that although intrinsically connected, these are separate processes that can be (mathematically) separated with complete datasets (i.e., those including cold hardiness and time to budbreak). The correlation between cold hardiness and deacclimation rate is spurious, as seen using a large dataset comprised of weekly evaluations of both for many different species (Kovaleski, 2022; Fig. **3b**,**c**). During fall and early winter, when cold hardiness has not reached its maximum, both cold hardiness and deacclimation rate increase, suggesting such correlation to be true. Once a maximum cold hardiness is reached for a given species {in about December for many species and maintained throughout winter [see Ferguson et al. (2011), Londo and Kovaleski (2017), North et al. (2021), Kovaleski (2022)]}, only the rate of deacclimation continues to increase in response to chilling accumulation. For some species that were evaluated throughout losing their field cold hardiness in early spring (*Acer rubrum* and *Cercis canadensis* in Fig. **3b**), we can observe that the rate of deacclimation can continue to increase even as the cold hardiness begins to decrease in the spring. Therefore, simply measuring the cold hardiness of buds does not say much about their dormancy state (or time to budbreak).

**Figure 3.**
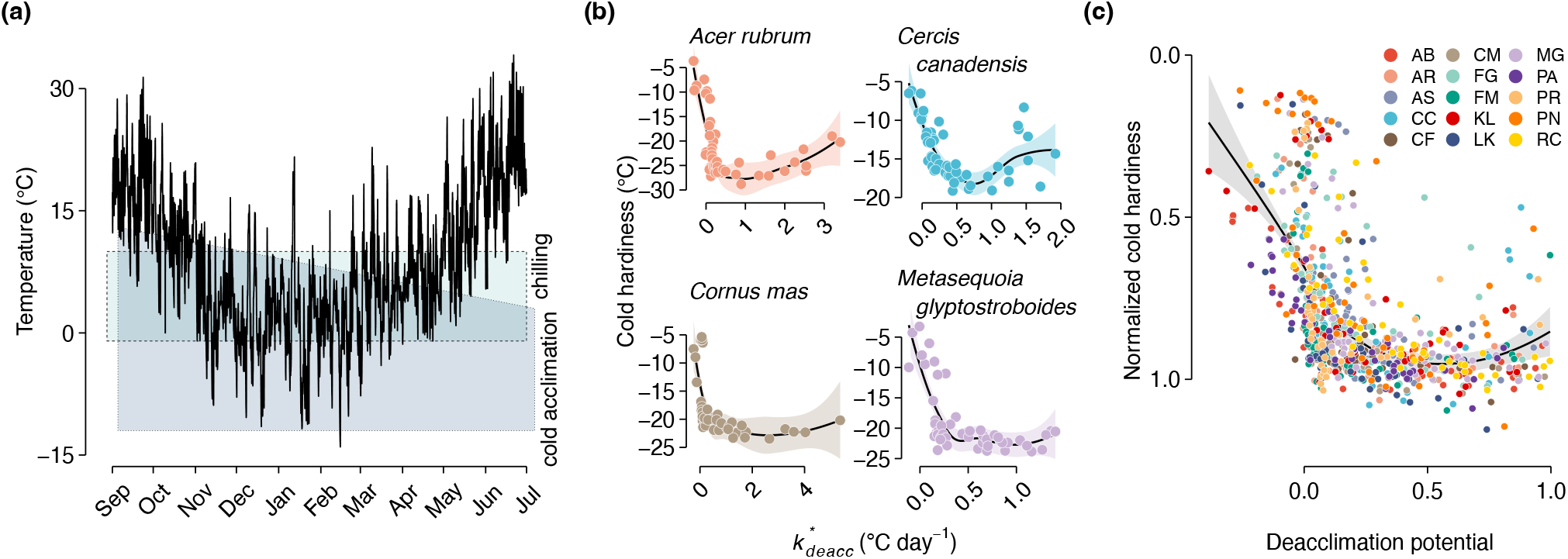
Cold hardiness and deacclimation rate are affected by low temperatures during the winter (data from Kovaleski, 2022). **(a)** Temperatures during 2019-2020 season in Boston, MA, USA, with overlayed effects of presumed temperatures eliciting chilling accumulation and cold acclimation responses. **(b)** Absolute values of cold hardiness and effective rate of deacclimation 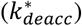 for four species of woody perennials. **(c)** Normalized values of cold hardiness and rate of deacclimation (dubbed deacclimation potential (Kovaleski et al., 2018)) for 15 species of woody perennials. AB – *Abies balsamea*; AR – *Acer rubrum*; AS – *Acer saccharum*; CC – *Cercis canadensis*; CF – *Cornus florida*; CM – *Cornus mas*; FG – *Fagus grandifolia*; FM – *Forsythia* ‘Meadowlark’; KL – *Kalmia latifolia*; LK – *Larix kaempferi*; MG – *Metasequoia glyptostroboides*; PA – *Picea abies*; PR – *Prunus armeniaca*; PN – *Prunus nigra*; RC – *Rhododendron calendulaceum*. Adapted from Kovaleski (2022).

Cold hardiness is therefore not a substitute for time to budbreak. To understand effects of chilling temperatures on dormancy, either (i) cold hardiness measurements must be attached to time to budbreak or (ii) deacclimation rates should be used. Considering budbreak is the culminating phenological event at the end of the dormant season, and useful in modeling, perhaps all three should be evaluated at once. Budbreak is an important phenological status when it comes to freeze risks of native vegetation and crops, which may benefit from protection. In addition, budbreak is easily observed, requiring no special equipment and thus allowing for field data collection by citizen science projects with much higher reach in terms of locations and number of individuals and species than would be possible if only done by scientists [e.g., Nature’s Notebook (Posthumus and Crimmins, 2011) within the USA National Phenology Network (www.usanpn.org), iNaturalist (www.inaturalist.org), and Pan European Phenological database (PEP 725; Templ et al., 2018)]. At the same time, however helpful extensive spring phenology datasets may be, thoughtful consideration must be made in experimental settings where detailed phenotyping is possible.

## Conclusions

Despite evidence presented here, some might still consider cold hardiness and dormancy to be a part of the same process. And differences in opinion are fertile ground for scientific innovation – something that appears needed for advances in dormancy research. Regardless, we believe to have shown direct and clear evidence here that future research in dormancy and spring phenology – be that in artificial or natural conditions – would benefit from including cold hardiness evaluation in their study designs. While we make a case for the effect of temperatures of chilling affecting cold hardiness, it is possible that any environmental effect that affects budbreak phenology {e.g., water (Hajek and Knapp, 2022), light [either photoperiod (Körner and Basler, 2010) or radiation (Vitasse et al., 2021)], and interactions (see Peaucelle et al., 2022)} may be doing so through affecting cold hardiness as well as dormancy. It is true that evaluation of cold hardiness of buds in dormancy studies may be more consequential in some species than others, but cold hardiness is, to our knowledge, always an intrinsic part of budbreak phenology. The full impact of acknowledging cold hardiness of buds may only be understood as more data is generated. We expect that this will not only help but will be crucial in elucidating aspects of dormancy mechanisms, as well as helping phenological modeling efforts.

Box 1. Current understanding of cold hardiness dynamics and its effect on time to budbreak

Most cold hardiness is lost upon budbreak (Lenz et al., 2013; Vitra et al., 2017; Kovaleski et al., 2018; Kovaleski, 2022), but the amount of cold hardiness and how it is lost can affect timing of budbreak. Examples here are based on extensive phenotyping of plants that use supercooling as a mechanism of cold hardiness, but we expect a similar dynamic for plants that use other mechanisms (Neuner et al., 2019; Villouta et al., 2020). Under forcing (i.e., exposure to warm temperatures and generally long days), supercooling ability is lost linearly, without changes in external morphology (Box **1a**) [but internal anatomical and morphological changes occur (Viherä-Aarnio et al., 2014; Xie et al., 2018; Kovaleski et al., 2019; Villouta et al., 2022)]. As growth resumes, the supercooling ability has been lost and concentration mostly drives cold hardiness of tissues. The minimum cold hardiness is thus observed at budbreak and early leafout (Chamberlain et al., 2019), when influx of water driving turgor of tissues leading to budbreak prior to influx of carbohydrates decreases concentration of tissues to a minimum. The relative alignment of these factors may vary based on a given definition of budbreak and/or morphological differences across species (Lancashire et al., 1991; Finn et al., 2007).

The supercooling ability is lost linearly relative to time under forcing conditions at a given temperature for many species (Kovaleski et al., 2018; Kovaleski, 2022; North et al., 2022). Here, this is illustrated conceptually using an orthogonal triangle. The time to budbreak is the base of the triangle, and the cold hardiness is the height of the triangle. The deacclimation rate (rate of cold hardiness loss) thus becomes the angle of the hypotenuse to the base of the triangle. Mathematically, these relations are represented by the following equation:

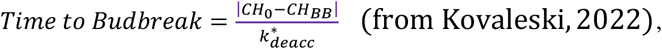

where *CH*_*0*_ is the initial cold hardiness, *CH*_*BB*_ is the cold hardiness at budbreak, 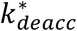 is the effective rate of deacclimation (a function of both temperature in which deacclimation is occurring and chill accumulation). Three scenarios are explored here where variations in cold hardiness and deacclimation rate affect timing of budbreak.

In the first example (Box **1b**), buds have the same initial cold hardiness, but deacclimate at different rates (red has higher rate than blue), thus leading to different times to budbreak (earlier for red than for blue). This may be caused by different levels of chill accumulation within the same species (or same genotype within a species), or different species at the same chill accumulation where one has inherently faster deacclimation rate. If these are the same genotype, the different rates mean that the buds are at different dormancy states.

In the second example (Box **1c**), buds have different initial cold hardiness (blue is more cold hardy than yellow), but deacclimate at the same rate. This could happen if buds are collected from the same genotype, at the same chill accumulation, but some buds were exposed to lower temperatures, leading to greater cold acclimation. A scenario where this could occur is buds collected in different locations, where one has lower minimum temperatures than the other. These being the same genotype, having the same deacclimation rate means the buds are at the same dormancy state, regardless of the difference in time to budbreak.

In the third example (Box **1d**), both initial cold hardiness and deacclimation rate are different (red is more cold hardy and has higher rate of deacclimation compared to yellow). Despite these differences, budbreak occurs at the same time. For the same genotype, this could be observed with less cold hardy buds in the fall, breaking bud in the same amount of time as buds collected in mid-winter which are more cold hardy, but lose that cold hardiness faster due to more chill accumulation. Although budbreak is happening at the same time, the buds are likely at different dormancy states.

In field conditions, bud cold hardiness follows a U-shaped pattern throughout the dormant period, while deacclimation rate increases in a sigmoid shape. When these two are combined, we find that observations of forcing experiments follow a certain pattern: time to budbreak increases slightly in fall where cold hardiness is increasing, but deacclimation rate has not yet significantly increased; this is followed by a period where cold hardiness stops increasing, and bud deacclimation rate rapidly increases, thus leading to decreases in time to budbreak; and finally, a period where deacclimation rate is no longer increasing (chilling is maximized), and cold hardiness starts to decrease.

These scenarios highlight the importance of a dormancy phenotype that integrates cold hardiness and deacclimation. Budbreak phenotyping alone overlooks important physiological differences associated with dormancy. In some cases, an integrated phenotype will support the interpretation of differing dormancy status based on budbreak but will enhance the extent of differences. In other cases, an integrated phenotype could greatly contradict interpretations of dormancy status based on budbreak.

**Fig. Box 1.**
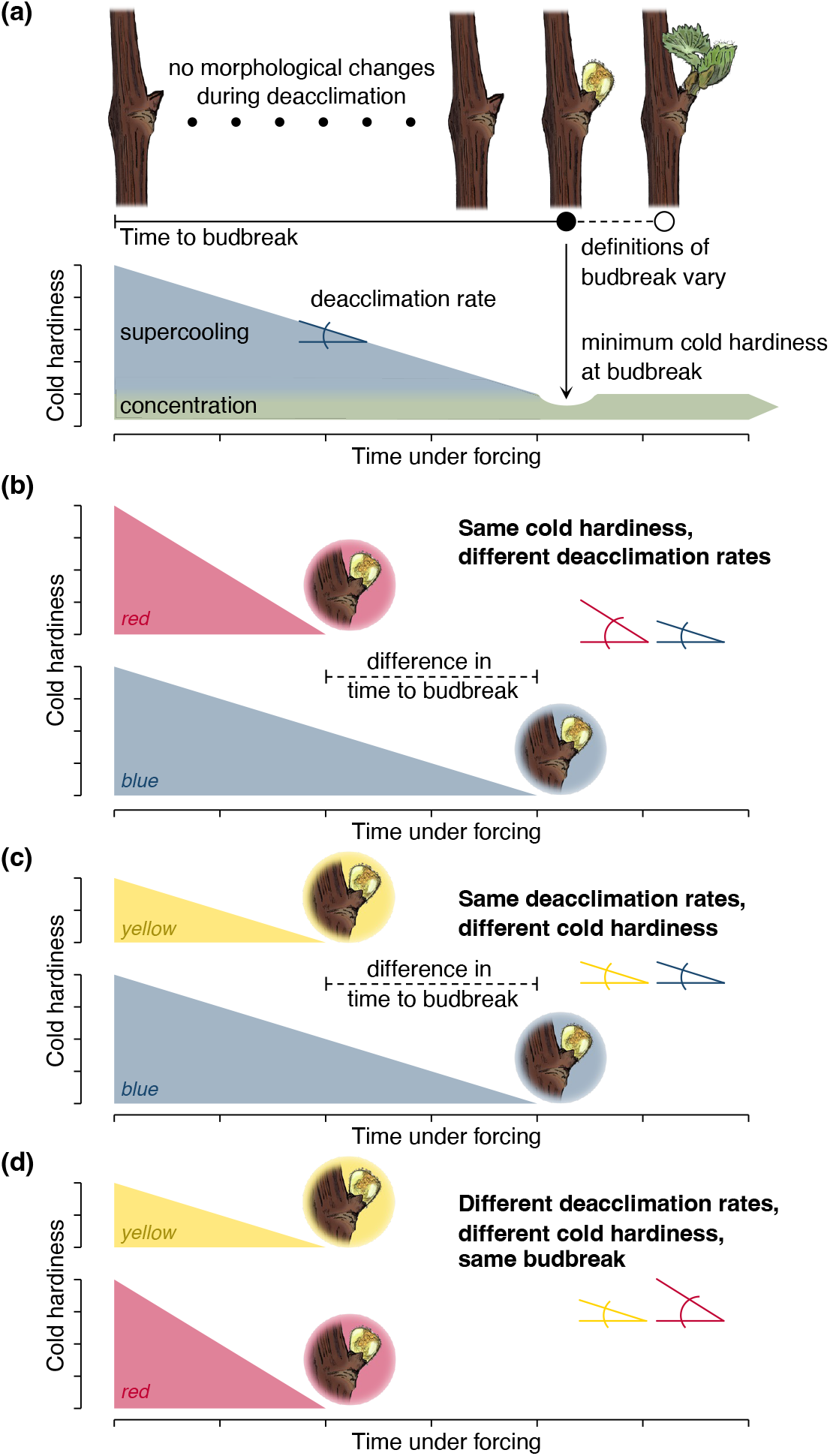
The time to budbreak is affected by degree of cold hardiness and deacclimation rate, as budbreak occurs once supercooling ability is lost. **(a)** In many species, cold hardiness of buds is determined by supercooling ability and cellular concentration. Upon budbreak, supercooling ability has been lost, and cold hardiness is minimal as concentration drops due to high turgor of tissues, with some being recovered as tissues mature. **(b–d)** Different scenarios are presented where: *initial cold hardiness* is the same for blue and red triangles, but lower for yellow triangles; *deacclimation rates* are the same for blue and yellow triangles, but greater for red triangles; and in combination resulting in *time to budbreak* being the same for yellow and red triangles, but greater for the blue triangle.

## Acknowledgements

We thank [names to be added post-review] for comments on drafts; the Arnold Arboretum of Harvard University for access to the living collections; and A. Atucha and the West Madison Agricultural Research Station for access to vineyards. Support for this research was provided by the Office of the Vice Chancellor for Research and Graduate Education at the University of Wisconsin–Madison with funding from the Wisconsin Alumni Research Foundation and the Putnam Fellowship Program of the Arnold Arboretum.

## Author contributions

MGN and APK designed research, performed research, analyzed data, wrote the manuscript.

## Notes

### Competing Interest Statement

The authors have declared no competing interest.

