## Supporting Information - Methods for "Time to budbreak is not enough: cold hardiness evaluation is necessary in dormancy and spring phenology studies"

### Materials and Methods

*Materials for Figure 1*. Cuttings with buds were collected from five interspecific hybrid *Vitis* cultivars [‘Brianna’ (BR), ‘Frontenac’ (FR), ‘La Crescent’ (LC), ‘Marquette’ (MQ), ‘Petite Pearl’ (PP)] in Madison, WI (43°03’37” N, 89°31’54” W) in winter of 2021-2022. Cuttings were incubated wrapped in a moist paper towel and sealed in a plastic bag in three different low temperature treatments: constant temperature (5 ºC), fluctuating temperature (–3.5 °C, 6.5 ºC, for 7h, 17h intervals daily), and field temperatures. After 2.5 months under treatment, cuttings from the constant and fluctuating treatments were reciprocally exchanged for 1 month of additional treatment. At approximately bi-weekly intervals, 15 to 30 buds from each cultivar and treatment were evaluated for cold hardiness using DTA (see *Cold hardiness evaluation* below). At the same time as the final cold hardiness evaluation, 15 cuttings from each cultivar and treatment were placed under forcing conditions to observe time to budbreak (see *Forcing assays* below).

*Materials for Figure 2a*. Cuttings with buds were collected from nine ornamental and forest species (*Acer platanoides*, *Acer rubrum*, *Acer saccharum*, *Cornus mas*, *Forsythia* ‘Meadowlark’, *Larix kaempferi*, *Metasequoia glyptostroboides*, *Picea abies*, *Prunus armeniaca*) in Boston, MA (42° 17’57”N, 71°07’22”W) on 25 November 2019. Cuttings were incubated upright in 2-in pots with proximal ends submerged in water in five different low temperature chilling treatments: four constant at –3 ºC, 4 ºC, 7 ºC, 11 ºC and one fluctuating at (–8 ºC, –3 ºC, 2ºC, –3 ºC for 6h intervals each; “–3±5 ºC”). Ten buds from each treatment and each species in each date were evaluated for cold hardiness using DTA at approximately weekly intervals (though not always 10 peaks were observed).

*Materials for Figure 2b*. Cuttings with buds were collected from eleven ornamental and forest species [*Abies balsamea* (AB), *Acer rubrum* (AR), *Acer saccharum* (AS), *Cercis canadensis* (CC), *Cornus florida* (CF), *Cornus mas* (CM), *Fagus grandifolia* (FG), *Larix kaempferi* (LK), *Metasequoia glyptostroboides* (MG), *Prunus armeniaca* (PR)] in Boston, MA (42° 17’57” N, 71°07’22” W) om 20 October 2020. Cuttings were incubated upright in 2-in pots with proximal ends submerged in water in low temperature treatments decreasing in –2 ºC steps every 10 days from 0 ºC to –6 ºC. Ten buds from each species in each date were evaluated for cold hardiness using DTA at approximately 2, 5, and 10 days post each temperature change (though not always 10 peaks were observed).

*Materials for Figure 3*. See "Woody species do not differ in dormancy progression: differences in time to budbreak due to forcing and cold hardiness" (Kovaleski, 2022; https://github.com/apkovaleski/dormancy_ArnoldArb).

*Data analysis*. All data were analyzed, and figures produced using R within R studio, and packages within [agricolae (De Mendiburu, 2009), dplyr (Wickham et al., 2021), ggbeeswarm (Clarke and Sherrill-Mix, 2017), ggh4x (van den Brand, 2021), ggplot2 (Wickham, 2016), patchwork (Pendersen, 2020), tidyr (Wickham and Girlich, 2022)].

*Figure 1b*: Simple linear regression was used to analyze cold hardiness and budbreak data in two separate models, one model with LTE as the response variable and the second model with days to budbreak as the response variable. Low temperature treatment (categorical variable for constant, fluctuating, field, con-flu, flu-con treatments) was used as the explanatory variable in both models. Means were then separated across treatments using Fisher's LSD at $\alpha = 0.05$.

*Figure 1c*: Simple linear regression using average time to budbreak as the response variable and average low temperature exotherm as the explanatory variable.

*Figure 2a*: Multiple linear regression was used to analyze acclimation (or lack of thereof) under low temperature treatments. A model was fit with average time to budbreak as the response variable. Temperature (categorical variable) and the interaction between temperature and time (continuous variable, i.e., days) were included as explanatory variables.

*Figure 2b*: Multiple linear regression was used to analyze acclimation (or lack of thereof) for different species under low temperature treatments. A model was fit with average low temperature exotherm as the response variable. Species (categorical variable) and the interaction between species and time (continuous variable, i.e., days) were included as explanatory variables.

*Forcing assays*. Cuttings were placed in cups under forcing conditions (22 °C, 16-h day/8-h night). Buds were observed for budbreak quasi-daily. Grapevine buds were considered to have broken when they reached stage 3 in the modified E-L scale (Coombe and Iland, 2005).

*Cold hardiness evaluation*. Differential thermal analysis (DTA) was conducted to estimate cold hardiness of buds by measuring their low temperature exotherms (Burke et al., 1976; Mills et al., 2006). The equipment used included acrylic trays fitted with thermoelectric modules (TEMs) to detect exothermic freezing reactions and a thermistor to monitor temperature in the trays. Trays were loaded in a Tenney JR programmable freezing chamber (Model TJR-A-F4T, Thermal Product Solutions) connected to a multimeter data acquisition system (Keithley 2700 for Figures 2 and 3 or DAQ6510 for Figure 1; Keithley Instruments). TEM voltage and thermistor resistance readings were collected via Keithley KickStart software (ver. 2.7.0). Voltage signals were examined graphically on Microsoft Excel to extract cold hardiness readings based on low temperature exotherm (LTE) peaks.

*Data availability*. Cold hardiness measurements, time to budbreak measurements, and code for analyses have been deposited in GitHub (https://github.com/apkovaleski/CH_to_budbreak) (Kovaleski, 2022b).
